# Novel 3,6-Dihydroxypicolinic Acid Decarboxylase Mediated Picolinic Acid Catabolism in *Alcaligenes faecalis* JQ135

**DOI:** 10.1101/457895

**Authors:** Qiu Jiguo, Zhang Yanting, Yao Shigang, Ren Hao, Qian Meng, Hong Qing, Lu Zhenmei, He Jian

## Abstract

*Alcaligenesfaecalis* strain JQ135 utilizes picolinic acid (PA) as sole carbon and nitrogen source for growth. In this study, we screened a 6-hydroxypicolinic acid (6HPA) degradation-deficient mutant through random transposon mutagenesis. The mutant hydroxylated 6HPA into an intermediate, identified as 3,6-dihydroxypicolinic acid (3,6DHPA) with no further degradation. A novel decarboxylase PicC was identified that was found to be responsible for the decarboxylation of 3,6DHPA to 2.5-dihydroxypyridine. Although, PicC belonged to amidohydrolase_2 family, it shows low similarity (<45%) when compared to other reported amidohydrolase_2 family decarboxylases. Moreover, PicC was found to form a monophyletic group in the phylogenetic tree constructed using PicC and related proteins. Further, the genetic deletion and complementation results demonstrated that *picC* was essential for PA degradation. The PicC was Zn^2+^-dependent non-oxidative decarboxylase that can specifically catalyze the irreversible decarboxylation of 3,6DHPA to 2.5-dihydroxypyridine. The *K*_m_ and *k*_cat_ towards 3,6DHPA were observed to be 13.44 μM and 4.77 s^-1^, respectively. Site-directed mutagenesis showed that His163 and His216 were essential for PicC activity.

**Importance:** Picolinic acid is a natural toxic pyridine derived from L-tryptophan metabolism and some aromatic compounds in mammalian and microbial cells. Microorganisms can degrade and utilize picolinic acid for their growth, and thus, a microbial degradation pathway of picolinic acid has been proposed. Picolinic acid is converted into 6-hydroxypicolinic acid, 3,6-dihydroxypicolinic acid, and 2,5-dihydroxypyridine in turn. However, there was no physiological and genetic validation for this pathway. This study demonstrated that 3,6DHPA was an intermediate in PA catabolism process and further identified and characterized a novel amidohydrolase_2 family decarboxylase PicC. It was also shown that PicC could catalyze the decarboxylation process of 3,6-dihydroxypicolinic acid into 2,5-dihydroxypyridine. This study provides a basis for understanding PA degradation pathway and the underlying molecular mechanism.

## Introduction

Decarboxylation is a fundamental process in nature (1, 2). A variety of organic compounds, including carbohydrates, fatty acids, aromatic compounds, and environmental xenobiotics, are involved in decarboxylation. The decarboxylase family can be subdivided into two groups based on the cofactor involved (1). Some enzymes require organic cofactors, such as flavin or NAD(P)^+^; while others utilize inorganic cofactors, such as Zn^2+^ or Mn^2+^. Recently, the amidohydrolase_2 family decarboxylases that use inorganic ions as cofactors, have been gaining more attentions (3-6). The enzymes in this family are usually involved in the catabolism of important natural compounds such as *α*-amino-*β*-carboxymuconate-ε-semialdehyde (ACMSD) (7, 8), *γ*-resorcylate (9), 2,3-dihydroxybenzoate (10), 2,5-dihydroxybenzoate (11), 3,4-dihydroxybenzoate (2), 4-hydroxybenzoate (12), 5-carboxyvanillate (13), vanillate (14), and 2-hydroxy-1-naphthoate (3). However, most of these studied compounds are benzene ring derivatives, while no decarboxylase has been studied that is involved in the catabolism of pyridine derivatives.

Picolinic acid (PA) is a typical C2-carboxylated pyridine derivate that is widely produce from physiological metabolism in mammalian and microbial cells (15). PA is a natural dead-end metabolite of L-tryptophan produced via kynurenine pathway in humans and other mammals (16-18). Moreover, it can be produced in other biological processes such as the microbial degradation of 2-aminophenol, catechol, and nitrobenzene (19-21). PA was found to be toxic and it inhibited the growth of normal rat kidney cells and T cell proliferation, thus, enhancing seizure activity in mice, and inducing cell death via apoptosis (22-25). PA cannot be metabolized by humans, thus gets excreted through urine or sweat (26). However, PA can be degraded by microorganisms in the natural environment (15). Numerous PA-degrading bacterial strains have been isolated including *Achromobacter* (27), *Aerococcus* (28), *Alcaligenes* (29), *Arthrobacter* (30), *Bacillus* (31), *Burkholderia* (32), or *Streptomyces* (33). The metabolic pathway of PA in microorganisms has been partially elucidated in previous studies (15, 28, 32) (Fig. 1). In other studies, the crude enzyme facilitating the conversion of PA to 6HPA has been preliminarily purified in *Arthrobacter picolinophilus* DSM 20665 and an unidentified gram-negative bacterium (designated as UGN strain) (30, 34). Nevertheless, the functional genes or enzymes involved in PA degradation has not been cloned or characterized yet.

**Fig. 1.**
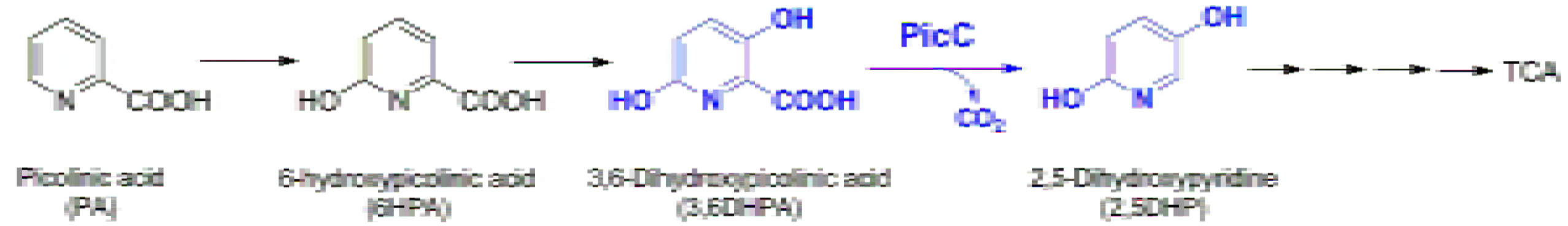
Proposed PA degradation pathway in *A. faecalis* JQ135. The 3,6DHPA and 2,5‐ DHP are shown in blue color. TCA, tricarboxylic acid cycle.

In our previous work, we demonstrated that *Alcaligenes faecalis* strain JQ135 utilizes PA as sole carbon, nitrogen, and an energy source and 6-hydroxypicolinic acid (6HPA) was the first intermediate of PA.(35). Further studies showed that *maiA* gene was essential for PA catabolism (36). In the present research, we reported the fully characterized intermediate compound, 3,6-dihydroxypicolinic acid (3,6DHPA) (Fig. 1). Further, a novel non-oxidative 3, 6-dihydroxypicolinic acid decarboxylase gene (*picC*) was cloned from *A. faecalis* strain JQ135, and the respective product was characterized.

## Results

### Transposon mutant and identification of the intermediate 3,6DHPA

A library of *A. faecalis* JQ135 mutants incapable of 6HPA utilization was constructed by random transposon mutagenesis. More than 30 mutants that could not grow on 6HPA were selected from approximately 10 000 clones and their ability to convert 6HPA was examined. The concentration of 6HPA was 1 mM and the inoculum of mutant was set at a final OD_600_ of 2.0. HPLC results showed that one mutant (designated as Mut-H4) could convert 6HPA into a new intermediate with no further degradation (Fig. 2). After liquid chromatography/time of flight-mass spectrometry (LC/TOF-MS) analysis, it was found that the molecular ion peak ([M+H]^+^) of this new intermediate was 156.0295 (Ion Formula, C_6_H_6_NO_4_^+^, calculated molecular weight 156.0297 with −3.2 ppm error), indicating that one oxygen atom was added to 6HPA (C_6_H_5_NO_3_). According to previously predicted PA degradation pathway, the intermediate is most likely to be 3,6DHPA (15, 31, 34). In the present study, 3,6DHPA was chemically synthesized and characterized by UV-visible, LC/TOF-MS, ^1^H NMR and ^13^C NMR spectroscopies (Fig. S1 and S2) and HPLC analysis showed that the retention time of the new intermediate was identical to that of the synthetic sample of 3,6DHPA (Fig. 2). Thus, this intermediate compound was identified as 3,6DHPA.

**Fig. 2.**
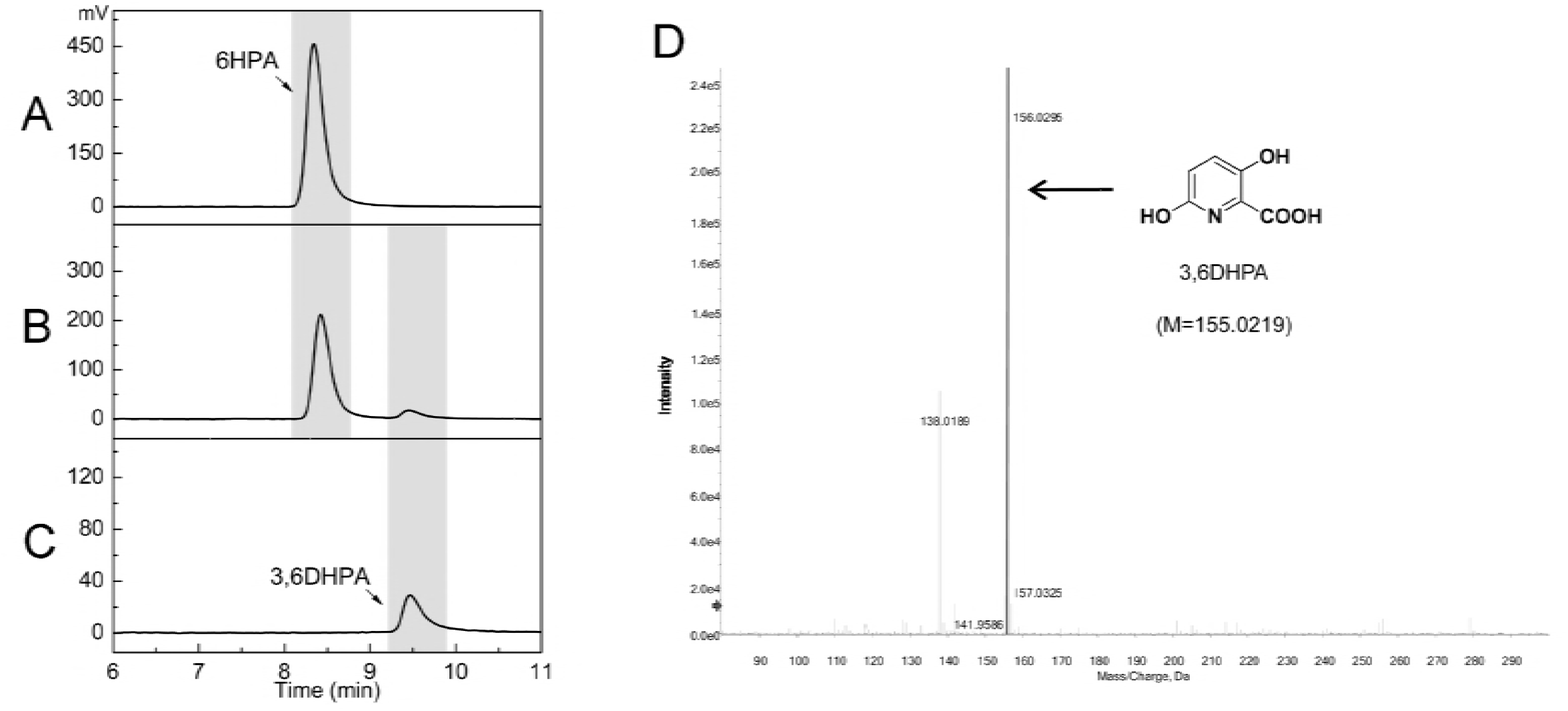
HPLC and LC/TOF-MS profiles of the conversion of 6HPA by mutant Mut-H4. (A and C) the authentic sample of 6HPA and 3,6DHPA, respectively. (B) The conversion of 6HPA into 3,6DHPA by mutant Mut-H4. The detection wavelength was set at 310 nm. (D) LC/TOF-MS spectra of 3,6DHPA produced in panel B.

### Screening of the 3,6DHPA decarboxylase gene

The transposon insertion site of mutant Mut-H4 was identified using the genomic walking method (37). The insertion site of the transposon was located in gene *AFA*_*15145* (genome position 3298929). Gene *AFA*_*15145* was a 972 bp length ORF starting with GTG. AFA_15145 exhibited the highest sequence similarities to several non-oxidative decarboxylases such as *γ*-resorcylate decarboxylase (*γ*-RSD, 45% identity) (9), 2,3-dihydroxybenzoate decarboxylase (2,3DHBD, 36% identity) (10), 5-carboxyvanillate decarboxylase (5CVD, 27% identity) (13), and hydroxynaphthoate decarboxylase (HndA, 22% identity) (3) (Fig. 3). All these decarboxylases belong to the amidohydrolase_2 family proteins (COG2159) that contain a triosephosphate isomerase (TIM)-barrel fold. Based on the phenotype of the mutant Mut-H4 and bioinformatics analysis, it was predicted that the *AFA*_*15145* (designated as *picC*) encoded the 3,6DHPA decarboxylase.

**Fig. 3.**
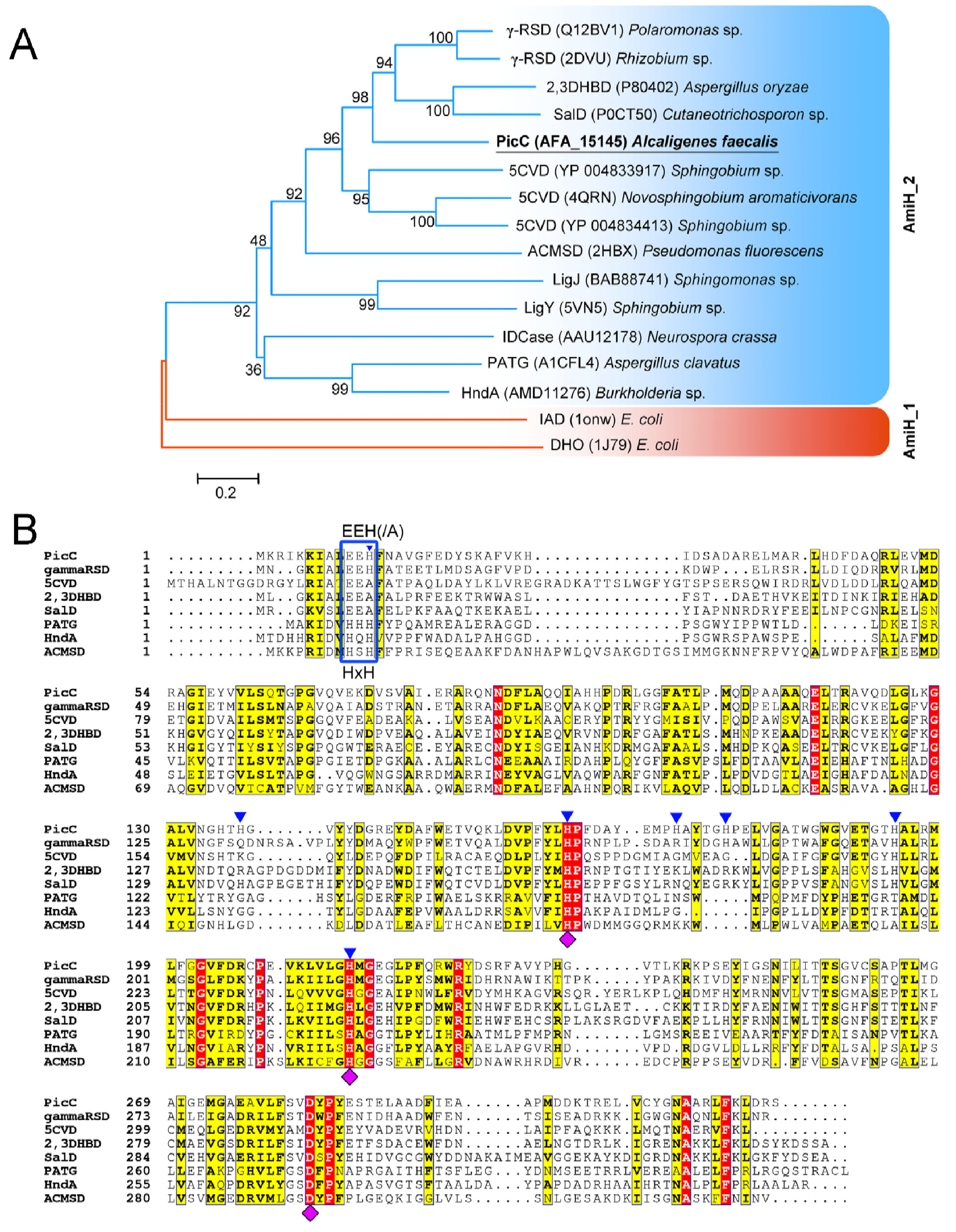
Amino acid sequence analysis of PicC. (A) Phylogenetic analysis of PicC and related decarboxylases. Each item was arranged in the following order: protein name, accession number, and strain. AmiH_1 and _2 are amidohydrolase_1 and _2 family, respectively. The phylogenetic tree was constructed using the neighbor-joining method (with a bootstrap of 1000) with software MEGA 6.0. The bar represents amino acid substitutions per site. (B) Multiple sequence alignment of PicC and seven decarboxylases. The predicted N-terminal motifs for Zn^2+^ binding are denoted by blue box. The other three Zn^2+^-binding sites are denoted by purple diamonds. The seven histidine residues for site-directed mutagenesis are denoted by blue triangles.

### Function identification of *picC* gene in PA degradation in *A. faecalis* JQ135

To confirm whether *picC* is involved in PA degradation, a *picC*-deleted mutant JQ135Δ*picC* was constructed. The mutant JQ135Δ*picC* lost the ability to grow on PA, 6HPA, or 3,6DHPA. The complementation strain, JQ135Δ*picC*/pBBR-*picC* completely restored the phenotype of growth on PA, 6HPA, and 3,6DHPA. These results showed that *picC* gene was essential for the degradation of PA in *A. faecalis* JQ135.

### The *picC* encodes 3,6DHPA decarboxylase

The recombinant PicC was overexpressed in *E. coli* BL21(DE3) cells containing the plasmid pET-*picC*. SDS/PAGE analysis showed the presence of an intense band, consistent with the 6 × His-tagged PicC (37 kDa) (Fig. 4). The degradation of 3,6DHPA by purified PicC was monitored spectrophotometrically. The maximum absorption was shifted from 340 nm (3,6DHPA) to 320 nm (2,5DHP). LC/TOF-MS analysis suggested that the molecular ion peak of the product was 112.0400 (M+H^+^), which was identical to that of 2,5DHP (36, 38). Further, the HPLC analysis showed that the retention time of the product was identical to that of the authentic sample of 2,5DHP. The 3,6DHPA was degraded completely with the formation of equal molar of 2,5DHP. Moreover, the PicC did not catalyze the reverse carboxylation of 2,5DHP in a reaction mixture containing NaHCO_3_.

**Fig. 4.**
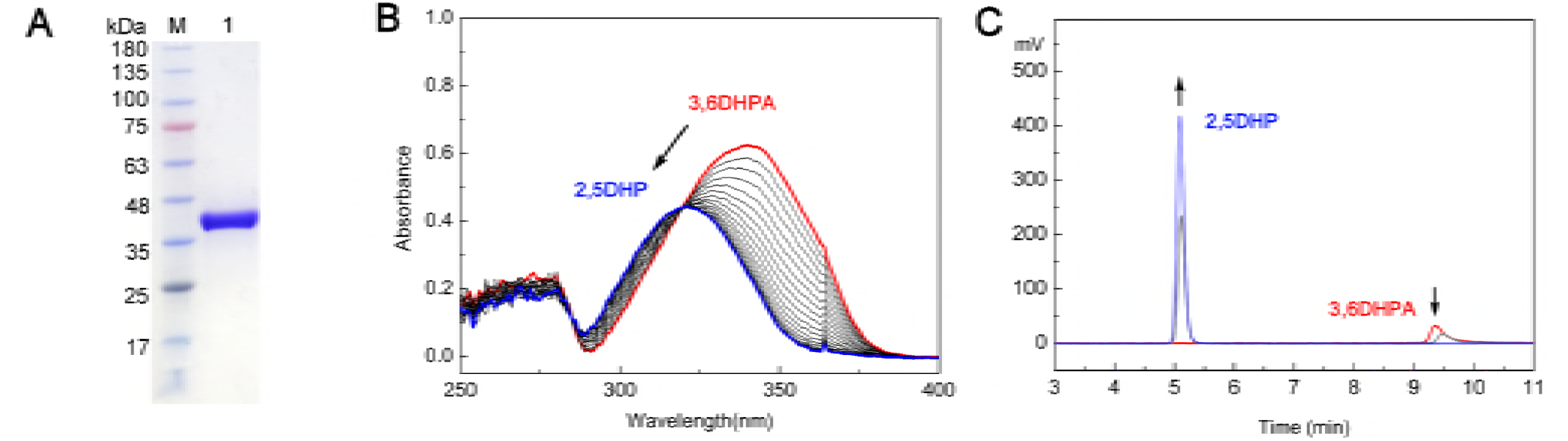
Characterization of PicC. (A) SDS-PAGE of purified PicC. Lane M, protein marker. Lane 1, purified PicC. (B) Spectrophotometric changes during transformation of 3,6DHPA by purified PicC. The reaction was initiated by adding 3,6DHPA. Spectra were recorded every 1 min. The arrow denotes the biotransformation of 3,6DHPA into 2,5DHP. (C) HPLC analysis of the transformation of 3,6DHPA into 2,5DHP by PicC. The detection wavelength was 310 nm.

### Biochemical properties of PicC

The recombinant PicC was highly active at pH 7.0 and 40°C (Fig. S3). The *K*_m_ and *k*_cat_ values for 3,6DHPA were found to be 13.44 μM and 4.77 s^-1^, respectively (Table 1). The enzyme was unstable at room temperature and could retain only 50% of initial activity when incubated at 30°C for 24 h. In addition, PicC could not convert the structural analogues of 3,6DHPA including 3-hydroxy-picolinic acid, gentisic acid, 2,3-dihydroxybenzoic acid, and 2,6-dihydroxybenzoic acid. This can be attributed to the substrate specificity of PicC towards 3,6DHPA.

**Table 1.** Kinetic constants of wild-type PicC and mutants.

| Enzyme | $K_m$<br>($\mu\text{M}$) | $V_{\max}$<br>($\mu\text{mol min}^{-1} \text{mg}^{-1}$) | $k_{\text{cat}}$<br>($\text{s}^{-1}$) | $k_{\text{cat}}/K_m$<br>($\text{s}^{-1} \text{mM}^{-1}$) |
| --- | --- | --- | --- | --- |
| PicC | $13.44 \pm 2.91$ | $7.73 \pm 0.05$ | 4.77 | 354.54 |
| PicC<sup>H12A</sup> | $53.06 \pm 4.05$ | $2.83 \pm 0.09$ | 1.75 | 32.94 |
| PicC<sup>H135A</sup> | $9.38 \pm 0.68$ | $6.14 \pm 0.11$ | 3.78 | 403.40 |
| PicC<sup>H163A</sup> | ND | ND | ND | ND |
| PicC<sup>H172A</sup> | $15.80 \pm 1.21$ | $3.33 \pm 0.08$ | 2.06 | 130.14 |
| PicC<sup>H177A</sup> | $110.81 \pm 14.70$ | $2.18 \pm 0.01$ | 1.35 | 12.14 |
| PicC<sup>H194A</sup> | $11.38 \pm 0.64$ | $5.15 \pm 0.07$ | 3.18 | 279.05 |
| PicC<sup>H216A</sup> | ND | ND | ND | ND |

ND: not detected

The effects of various metal ions and inhibitors, on decarboxylase activity are presented in Table 2. PicC activity was not affected by metal ions such as Ca^2+^, Cd^2+^, Co^2+^, Fe^3+^, Mg^2+^, Mn^2+^, and Zn^2+^ but was strongly inhibited by Ag^+^, Co^2+^, and Hg^2+^ ions. In addition, several inhibitors such as EDTA, 8-hydroxy-quinoline-5-sulfonic acid (8-HQSA, a zinc metal specific inhibitor), phenylmethylsulfonyl fluoride (PMSF, serine and cysteine specific inhibitor), and sodium iodoacetate (cysteine specific inhibitor) showed relatively low effects on PicC activity. However, diethylpyrocarbonate (DEPC), a histidine residue modifier, strongly inhibited PicC activity, indicating the presence of active-site histidine residues.

**Table 2.**
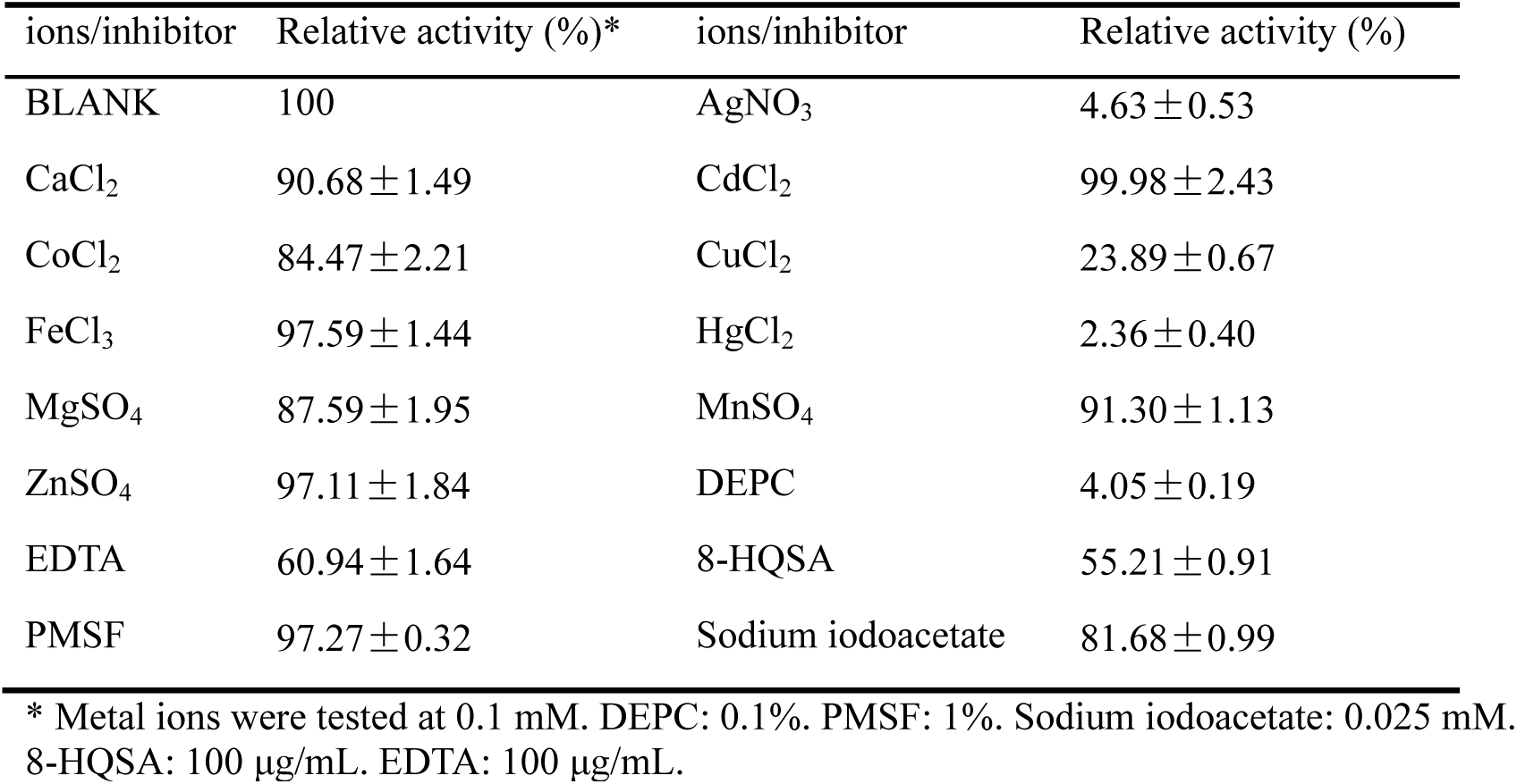
Effect of metal ions and inhibitors on PicC activity.

| ions/inhibitor | Relative activity (%)* | ions/inhibitor | Relative activity (%) |
| --- | --- | --- | --- |
| BLANK | 100 | AgNO<sub>3</sub> | 4.63 ± 0.53 |
| CaCl<sub>2</sub> | 90.68 ± 1.49 | CdCl<sub>2</sub> | 99.98 ± 2.43 |
| CoCl<sub>2</sub> | 84.47 ± 2.21 | CuCl<sub>2</sub> | 23.89 ± 0.67 |
| FeCl<sub>3</sub> | 97.59 ± 1.44 | HgCl<sub>2</sub> | 2.36 ± 0.40 |
| MgSO<sub>4</sub> | 87.59 ± 1.95 | MnSO<sub>4</sub> | 91.30 ± 1.13 |
| ZnSO<sub>4</sub> | 97.11 ± 1.84 | DEPC | 4.05 ± 0.19 |
| EDTA | 60.94 ± 1.64 | 8-HQSA | 55.21 ± 0.91 |
| PMSF | 97.27 ± 0.32 | Sodium iodoacetate | 81.68 ± 0.99 |

\* Metal ions were tested at 0.1 mM. DEPC: 0.1%. PMSF: 1%. Sodium iodoacetate: 0.025 mM. 8-HQSA: 100 µg/mL. EDTA: 100 µg/mL.

Further, absorption spectroscopy analysis revealed the presence of Zn^2+^ at 0.85±0.1 mol per mol of protein that is similar to several other non-oxidative decarboxylases of the amidohydrolase_2 superfamily (1, 3). The moderate effects, exhibited by additional Zn^2+^ or EDTA in the reaction system, indicated the presence of Zn^2+^ in the center of PicC.

### Site-directed mutagenesis

In order to assess their roles in the function of PicC, seven histidine residues (H12, H135, H163, H172, H177, H194, and H216) were substituted with Ala residue through site-directed mutagenesis (Fig. 3B). The seven PicC mutants obtained were expressed and purified and their activities were determined (Fig. S4; Table 1). PicC^H135A^ showed a slight increase in decarboxylase activity, while PicC^H172A^ and PicC^H194A^ showed a slight reduction in decarboxylase activities (10%∼50%) and PicC^H12A^ and PicC^H177A^ strongly reduced the decarboxylase activities (>90%). Moreover, the mutant proteins PicC^H163A^ and PicC^H216A^ completely lost their decarboxylase activities.

## Discussion

PA is a natural and toxic mono-carboxylated pyridine derivative. The studies on the microbial degradation mechanism of PA began 50 years ago (28). A partial catabolic pathway of PA has been proposed (Fig. 1): PA was dehydrogenated to 6HPA, and then gets converted into 3,6DHPA via hydroxylation leading to the decarboxylation of 3,6DHPA into 2,5DHP. The intermediate, 6HPA has been substantially identified in most strains including *Aerococcus* sp. (28), *Alcaligenes faecalis* DSM 6269 (29), *Arthrobacter picolinophilus* DSM 20665 (30, 39), *Burkholderia* sp. ZD1 (32), *Streptomyces* sp. Z2 (33), and the UGN strain (34). Further, another intermediate compound, 2,5DHP has been detected in the media during PA degradation in few strains (32, 34). However, the intermediate 3,6DHPA, a key link between 6HPA and 2,5DHP, was hardly detectable. This could be most attributed to its immediate degradation before excreting out of the cells. Previously, 3,6DHPA has only been theoretically proposed in *Bacillus* sp. (31) and the UGN strain (34). In this study, to the best of our knowledge, we demonstrated the chemical properties (UV-visible, LC/TOF-MS, ^1^H NMR and ^13^C NMR spectroscopies) of 3,6DHPA for the first time (Fig. S2) and detected it in the media using the transposon mutant strain, thus confirming that 3,6DHPA is a catabolic intermediate of PA.

Some previous studies have attempted to identify the genes and enzymes involved in PA degradation, such as the PA dehydrogenase in *Arthrobacter* (39) and 2,5DHP dioxygenase in UGN strain (34). However, their amino acid sequences and respective coding genes remained unknown, with no biochemical, physiological or genetic evidence to explain the decarboxylation of 3,6DHPA to 2,5DHP. In this context, we cloned a decarboxylase gene *picC* through random transposon mutagenesis, and ascertained that PicC was responsible for 3,6DHPA decarboxylation to form 2,5DHP. We found that PicC shared homology with the amidohydrolase_2 family proteins and contained the conserved triosephosphate isomerase (TIM)-barrel fold of amidohydrolase_2 family (Fig. 3). It has been previously reported that the amidohydrolase_2 family protein (PF04909) catalyzes the decarboxylation reaction (C-C bond) of several benzene derivatives, whereas the amidohydrolase_1 family protein (PF01979) catalyzes the hydrolytic reactions (C-N, C-Cl, or C-P bond) (4). The ACSMD was the first member of amidohydrolase_2 family to be reported (8) followed by other members, including *γ*-RSD (9), 2,3DHBD (10), 5CVD (13), and HndA (3). A phylogenetic tree of PicC and related proteins showed that PicC was clustered with amidohydrolase_2 but not amidohydrolase_1 family proteins (Fig. 3). However, the identities between PicC and reported decarboxylases were low (less than 45%), and PicC formed a separate branch in the phylogenetic tree (Fig. 3). In addition, PicC was found to be specific toward its substrate 3,6DHPA. Thus, it can be concluded that PicC could be a novel amidohydrolase_2 family decarboxylase.

The amidohydrolase_2 family proteins contain a few conserved amino acid residues, which are usually the active sites. In ACSMD, the His177, His228, and D294 were the Zn^2^+-binding sites (4). These three residues have been found in all reported amidohydrolase_2 family proteins including PicC (His163, His216, and D283) (Fig. 3B). In addition, the results of site-directed mutagenesis of PicC confirmed that H163 and H216 also played essential roles in PicC-mediated catalysis. Another Zn^2^+-binding motif 'HxH' has been found in the N-terminal of ACSMD (4) and HndA (3), whereas this motif was replaced by 'EEH' in *γ*-RSD (9) or 'EEA' in 5CVD (13) (Fig. 3B). In PicC, the corresponding motif has been found to be similar to that of *γ*-RSD. After substituting His12 by Ala, the resultant enzyme PicC^H12A^ still exhibited 10% activity, suggesting a variation in the third residue of this motif. In addition, site-directed mutagenesis results demonstrated that several other histidine residues His172, His177, and His194 were important for PicC activity.

In conclusion, this study revealed that 3,6DHPA was a catabolic intermediate in PA degradation by bacteria. The 3,6DHPA decarboxylase (PicC) was identified and characterized. To the best of our knowledge, PicC is also the first non-oxidative decarboxylase belonging to the amidohydrolase_2 family that catalyzes the irreversible decarboxylation of pyridine derivative. This study will expand our understanding of the bacterial degradation mechanisms of pyridine derivatives.

## Materials And Methods

### Chemicals

PA, 6HPA, and 2,5DHP were purchased from J&K Scientific Ltd. (Shanghai, China). EDTA, 8-HQSA, PMSF, sodium iodoacetate, DEPC, and other reagents of analytical grade were purchased from Sangon Biotech Co., Ltd. (Shanghai, China). 3,6DHPA was chemically synthesized (detailed in the supplemental materials). The structure of 3,6DHPA was confirmed by UV-visible, LC/TOF-MS, and NMR spectroscopy (Fig. S1 and S2).

### Strains, plasmids, and primers

All bacterial strains and plasmids used in this study are listed in Table 3. *Alcaligenes faecalis* JQ135 (CCTCC M 2015812) is the wild-type PA-degrading strain (35). *E. coli* DH5α was used as the host for the construction of plasmids. *E. coli* BL21(DE3) was used to over-express the proteins. Bacteria were cultivated in LB medium at 37°C (*E. coli*) or 30°C (*Alcaligenes* and their derivatives). Antibiotics were added at the following concentrations (as required): chloramphenicol (Cm), 34 mg/L; gentamicin (Gm), 50 mg/L; kanamycin (Km), 50 mg/L; and streptomycin (Str), 50 mg/L. Primer synthesis and the sequencing of PCR products or plasmids were performed by Genscript Biotech (Nanjing, China) (40). The primers used in this study are listed in Table 4.

**Table 3.**
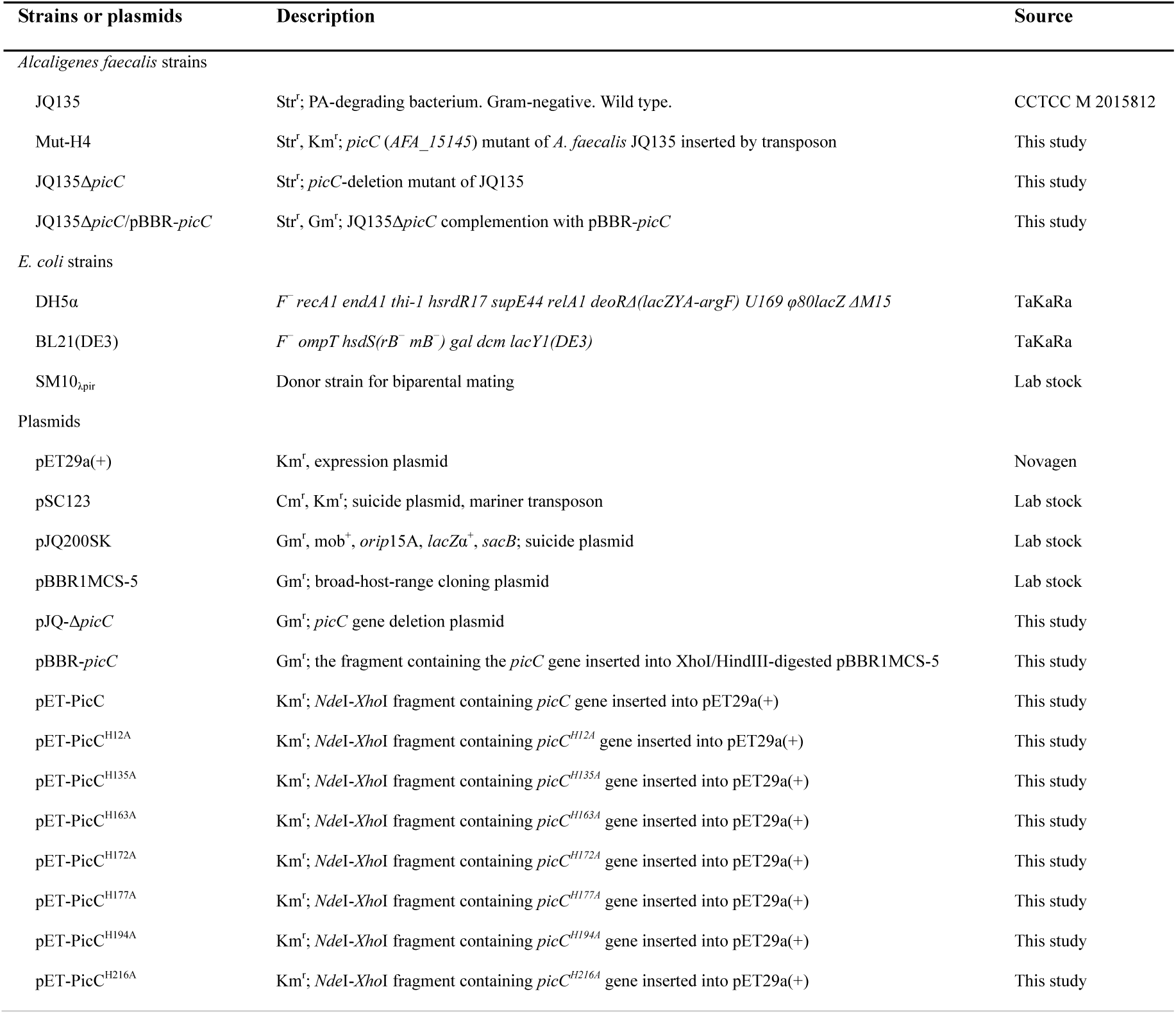
Strains and plasmids used in this study.

**Table 4.**
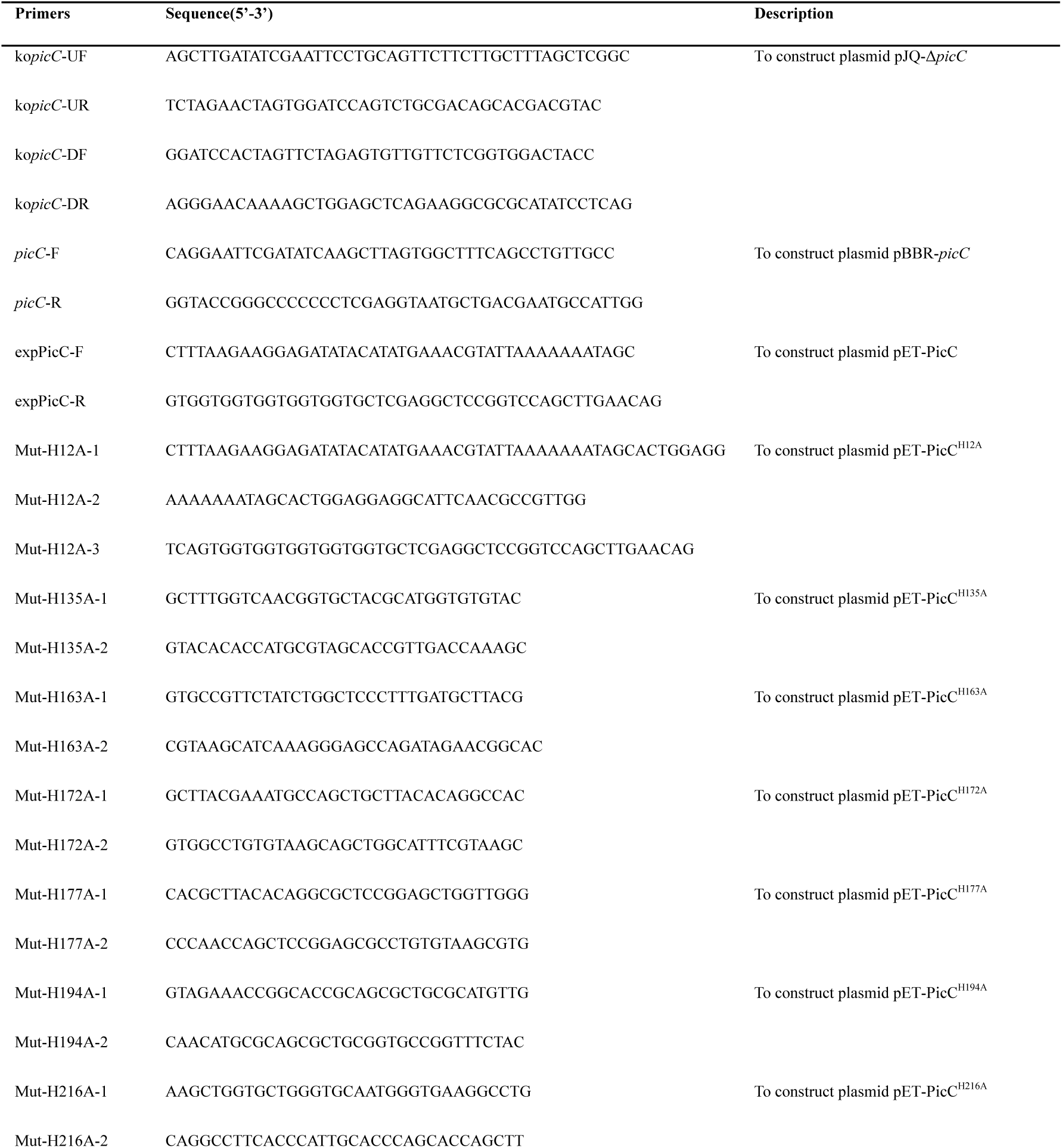
Primers used in this study.

### Transposon mutagenesis, mutant screening, and gene cloning

A transposon mutant library of *A. faecalis* JQ135 was constructed using the transposon-based plasmid pSC123 (kanamycin-resistance gene) as described previously (35). In this study, the PA was replaced by its intermediate 6HPA. Mutants that could not utilize 6HPA as the sole carbon source were selected. The flanking sequences of the transposon in the mutants were amplified using the DNA walking method (37). The amplified PCR products were sequenced and analyzed. The insertion sites were confirmed by comparison with the genome sequence of *A. faecalis* JQ135.

### Gene knockout and genetic complementation of *A. faecalis* JQ135

The genes or DNA fragments from *A. faecalis* JQ135 were were amplified by PCR using corresponding primers (Table 4). The fusion of DNA fragments and cut plasmids was carried out using the ClonExpress MultiS One Step Cloning Kit (Vazyme Biotech Co.,Ltd, Nanjing, China). Gene deletion mutant of the *picC* in *A. faecalis* JQ135 was constructed using a two-step homogenetic recombination method with the suicide plasmid pJQ200SK (41). Two homologous recombination-directing sequences were amplified using primers, ko*picC*-UF/ko*picC*-UR and ko*picC*-DF/ko*picC*-DR, respectively. Two PCR fragments were subsequently ligated into SacI/PstI-digested pJQ200SK generating pJQ-Δ*picC*. The *pJQ*-*ΔpicC* plasmid was then introduced into *A. faecalis* JQ135 cells. The single-crossover mutants were screened on a LB plate containing Str and Gm. The gentamicin-resistant strains were then subjected to repeated cultivation in LB medium containing 10% sucrose with no gentamicin. The double-crossover mutants that lost their plasmid backbone and were sensitive to gentamicin, were selected on LB Str plates. Deletion of the *picC* gene was confirmed by PCR. This procedure resulted in the construction of the deletion mutant strain JQ135Δ*picC*.

Knockout mutants were complemented as follows. The intact *picC* gene was amplified using the primers *picC*-F and *picC*-R, and then ligated with the Xhol/Hindlll-digested pBBR1-MCS5, generating pBBR-*picC*. The pBBR-*picC* was then transferred into the mutant strain JQ135Δ*picC* to generate the complemented strain JQ135Δ*picC*/pBBR-*picC*.

### Expression and Purification of the His-tagged PicC and its mutations

For the over-expression of *picC* gene in *E. coli* BL21(DE3), the complete ORF without the stop codon (genome position 3298274-3299242) were amplified using genomic DNA of strain JQ135 and inserted into the Ndel/Xhol-digested plasmid pET29a(+), resulting in the plasmid pET-PicC. *E. coli* BL21(DE3) cells (containing pET-PicC) were initiated by the addition of 0.3 mM IPTG when the optical density of the culture (OD_600_) reached 0.5-0.8 and was incubated for an additional 12 h at 16°C. Cells were harvested by centrifugation at 4 °C, sonicated, and then centrifuged again to remove cell debris. The supernatant was used for recombinant protein purification using Ni-NTA agarose column (Sangon, Shanghai, China). The purified 6 × His-tagged protein were then analyzed using 12.5% SDS-PAGE. The protein concentrations were determined using the Bradford method (42).

For site-directed mutagenesis of PicC, the *picC* fragments were amplified from plasmid pET-PicC through overlap PCR using the primers carrying point mutations (Table 4). Amplified fragments were fused into plasmid pET29a(+), resulting in pET-PicC^H12A^, pET-PicC^H135A^, pET-PicC^H163A^, pET-PicC^H172A^, pET-PicC^H177A^, pET-PicC^H194A^, and pET-PicC^H216A^. Resultant constructs were confirmed by sequencing. The expression and purification of the mutations were performed as described in the section above.

### Enzymatic assays of 3,6DHPA decarboxylase

For the decarboxylase activity, enzyme reaction mixture contained 50 mM PBS (pH 7.0), 0.3 mM 3,6DHPA, and 5 μg purified PicC (in 1 mL) and incubated at 40°C. The enzymatic activities were determined spectrophotometrically by the disappearance of 3,6DHPA at 360 nm (ε=4.4 cm^-1^ mM^-1^). To determine the effect of one condition, other conditions were kept at fixed concentration of the standard reaction. The optimum pH of the PicC protein was determined using various buffers such as 50 mM citric acid-sodium citrate (pH 4 to 6), 50 mM KH_2_PO_4_-K_2_HPO_4_ (pH 6 to 8), and 50 mM glycine-NaOH (pH 8.0 to 9.8) at 40°C. The optimum temperature of the PicC protein was determined to be in between 10°C to 50°C in PBS (pH 7.0). Purified PicC was pre-incubated with various metal ions and inhibitors at 4°C for 30 min to study their effects on the enzyme. The activity was expressed as a percentage of the activity obtained in the absence of the added compound. To determine the kinetic constants for 3,6DHPA, a range of 3,6DHPA concentrations (2 to 150 μM) were used. The values were calculated through non-linear regression fitting to the Michaelis-Menten equation. One unit of the activity was defined as the amount of enzyme that catalyzed 1 μmol of 3,6DHPA in 1 min.

The measurement of carboxylase activity of PicC was similar with a previous study (43). The reaction mixture contained 50 mM PBS (pH 7.0), 0.3 mM 2,5DHP, 5.0 mM NaHCO_3_ and 5 μg purified PicC in 1 mL mixture at 40 °C.

### Analytical methods

The UV-VIS spectra was observed by a UV2450 spectrophotometer (Shimadzu). The determination of PA and 6HPA, 3,6DHPA, and 2,5DHP concentrations were performed by HPLC analysis on a Shimadzu AD20 system equipped with a Phecda C18 reversed phase column (250 mm × 4.60 mm, 5 μm). The concentrations of the compounds were calculated using standard samples. The mobile phase was consisted of methanol : water : formic acid (12.5:87.5:0.2, v/v/v) at a flow rate of 0.6 mL/min, at 30 °C. LC/TOF-MS analysis was performed in a TripleTOF 5600 (AB SCIEX) mass spectrometer, as described previously (44). The Zn^2+^ concentration in the PicC protein was analyzed using inductively coupled plasma optical emission spectrometry (ICP-OES) according to a previous study (45).

### Nucleotide sequence accession numbers

The PicC and the complete genome sequence of *A. faecalis* JQ135 have been deposited in the GenBank database under accession numbers <u>ARS01287</u> and <u>CP021641</u>, respectively.

## Acknowledgments

We thank Dr. Chensi Shen (Donghua University) for the help of 3,6DHPA synthesis and MogoEdit Co. for providing linguistic assistance during the preparation of the manuscript.

This work was supported by the State's Key Project of Research and Development Plan (2016YFD0801102), the National Natural Science Foundation of China (Nos. 41630637, 31870092, and 31770117).

